# High performers demonstrate greater neural synchrony than low performers across behavioral domains

**DOI:** 10.1101/2023.06.22.546173

**Authors:** Taylor A. Chamberlain, Anna Corriveau, Hayoung Song, Young Hye Kwon, Kwangsun Yoo, Marvin M. Chun, Monica D. Rosenberg

## Abstract

Heterogeneity in brain activity gives rise to heterogeneity in behavior, which in turn comprises our distinctive characteristics as individuals. Studying the path from brain to behavior, however, often requires making assumptions about how similarity in behavior scales with similarity in brain activity. Here, we expand upon recent work which proposes a theoretical framework for testing the validity of such assumptions. Using intersubject representational similarity analysis in two independent movie-watching fMRI datasets, we probe how brain-behavior relationships vary as a function of behavioral domain and participant sample. We find evidence that, in some cases, the neural similarity of two individuals is not correlated with behavioral similarity. Rather, individuals with higher behavioral scores are more similar to other high scorers whereas individuals with lower behavioral scores are dissimilar from everyone else. Ultimately, our findings motivate a more extensive investigation of both the structure of brain-behavior relationships and the tacit assumption that people who behave similarly will demonstrate shared patterns of brain activity.

## Introduction

Individuals are amalgamations of their unique traits and abilities, and a key aim of cognitive neuroscience is uncovering how these personal characteristics arise from brain activity. But what is the nature of the relationship between individual differences in behavior and individual differences in brain activity?

If this question seems too theoretical to inform the day-to-day aspects of research, it is worth considering that often we answer it implicitly. Specifically, in modeling behavior, we frequently assume that people who behave similarly, as measured by similar scores on a behavioral metric, will demonstrate some shared pattern of brain activity. For instance, we might examine brain activity of participants with depression, expecting that depressed participants will exhibit more or less activity (or functional connectivity) in a given region compared to controls (Greicius et al., 2007; Siegle et al., 2002). Another popular method of relating patterns of brain activity to behavior is regression analysis. If we are interested in individual differences in sustained attention, we may opt to use regression to identify functional connectivity edges that are associated with better or worse attentional performance (Rosenberg et al., 2016; Yoo et al., 2022).

The idea that people who are similar in some regard will share some aspect of brain function is intuitive. But is it accurate? Returning to the example of depression research, suppose the patient sample composed of two Diagnostic and Statistical Manual of Mental Disorders subtypes, for instance melancholic and anxious (Musil et al., 2017). It is possible the melancholic subtype is characterized by less activation in the region of interest while the anxious subtype is associated with greater activation (or vice versa). Collapsing across patients could cause this effect to sum to zero, possibly leading to an average activation close to controls. In this way, by assuming that patients will appear similar in terms of brain activity, we run the risk of concluding that there is no significant difference between patients and controls and missing the underlying relationship entirely.

Indeed, several studies provide evidence that occasionally behavioral similarity (in this example, similar clinical symptoms) accompanies *less* similar brain activity. For instance, work shows that participants with autism spectrum disorder exhibit lower neural synchrony during movie-watching compared to controls (Hasson et al., 2009; Lyons et al., 2020; Salmi et al., 2013), as well as greater variability in inter-subject functional correlation states (Bolton et al., 2020). Intersubject correlation during movie-watching is also decreased among individuals with attention-deficit/hyperactivity disorder compared to controls (Salmi et al., 2020). Outside of intersubject correlation as a measure of variability, a study of individuals with autism spectrum disorder demonstrated greater inter-individual variability in fMRI activity (as measured by correlational distance between vectorized beta values) during a spatial working memory task (Hawco et al., 2020). Similarly, other work has shown greater variability in fMRI activity in patients with schizophrenia (Gallucci et al., 2022; Maïza et al., 2010).

Similar behavior, therefore, does not always imply similar brain activity. When should we expect this to be the case? Recent work by Finn and colleagues proposes a paradigm that can help elucidate the complex relationship between brain and behavioral similarity by empirically testing different models of brain-behavior relationships (Finn et al., 2020). The first model, the *Nearest Neighbors* model, operationalizes the assumption that those who score similarly on some behavioral metric will appear similar in brain activity. Studies testing for a linear brain-behavior relationship assume the *Nearest Neighbors* model. Alternatively, the second model, the *Anna Karenina (AnnaK)* model is named after the opening line of the novel *Anna Karenina*, stating “Happy families are all alike; every unhappy family is unhappy in its own way.” The *AnnaK* model proposes that one end of a behavioral spectrum is associated with greater variability in brain activity. They found that a measure of working memory showed an *AnnaK*-style relationship, where high scorers displayed greater neural synchrony with each other, whereas low-scoring participants were dissimilar to both low and high scorers.

Here we replicate and extend this finding in two independent fMRI datasets in which youth or adult participants watched movies during scanning. In doing so, we can determine to what extent the best model linking brain and behavioral similarity depends on the characteristics of the sample in question or on the specific behavior under investigation. Additionally, inclusion of a sample of children and adolescents specifically allows us to test if brain-behavior relationships look significantly different in development than they do in adulthood. Recent work found that a relationship between depression symptoms and intersubject correlation during movie watching emerges in adolescence, suggesting that brain-behavior relationships may vary as a function of age (Gruskin et al., 2020). Ultimately, across both the developmental and young adult sample, we find significant evidence for the *AnnaK* model. The *AnnaK* model fit our data across five measures: a cognitive function score, a language function score, two attention tasks and one working memory task. In contrast, the *Nearest Neighbors* only explains significant variance in neural similarity for a subset of measures where we find support for the *AnnaK*. As a whole, our findings illustrate that the neural similarity depends not only whether two individuals resemble each other behaviorally, but also on one’s position on a behavioral scale.

## Methods

### Dataset 1: Healthy Brain Network

#### Participants

To investigate how similarity in brain activity relates to behavioral similarity in development, we analyzed data from the Healthy Brain Network Biobank (Alexander et al., 2017). This project is collecting data from children and adolescents aged 5-21 years with a diversity of clinical concerns (the majority of the sample having one or more clinical diagnosis). The Healthy Brain Network project was approved by the Chesapeake Institutional Review Board. Secondary data analysis was approved by the University of Chicago Institutional Review Board. Of interest to our current analysis, the dataset includes one functional MRI (fMRI) run collected while participants watch an emotionally evocative video, specifically a 10-minute clip from the movie *Despicable Me*. In addition to fMRI data, this sample also contains a large assortment behavioral and phenotypic measures, including psychiatric and learning assessments, and questionnaires pertaining to environmental and lifestyle factors. We analyze a subset of these measures in the present study (see *Behavioral data* for details).

To determine our final study sample, we first downloaded all behavioral data available as of March 8, 2022 and all available neuroimaging data collected at sites Rutgers and Citibank Biomedical Imaging Center from Healthy Brain Network releases 1-8. Of the participants with downloaded data (*n*=2131), we subset the sample to those who had complete *Despicable Me* fMRI data and acceptable motion during this scan (defined as maximum head displacement <3 mm and mean framewise displacement <.15 mm) (*n*= 529). Of participants with a usable *Despicable Me* scan, we retained only participants who had behavioral data and whose anatomical scan passed visual quality control inspection (*n*=480).

From this point, behavioral measures available in 90% of the sample were retained. Of the remaining variables, two raters from our lab selected measures pertaining to five domains of interest (Cognitive, Attention, Social, Emotional, and Language) and grouped these measures by domain. Finally, participants missing any measure from any of the five domains were excluded resulting in a final sample of *n*=363 participants (142 F, 221 M; mean age = 12.25± 3.4 years, range = 6–21 years).

#### Behavioral data

To assess relationships between behavioral similarity and neural synchrony, we took advantage of the large variety of phenotypic and behavioral data provided by the Healthy Brain Network. The final measures comprising our five domains of interest (Cognitive, Attention, Social, Emotional, and Language) included self-report questionnaires, parent-completed questionnaires, and cognitive assessments such as the NIH toolbox (Hodes et al., 2013). For a complete list of measures included, please see Figure 2. Rather than analyze each measure of interest independently, we ran principal component analyses (PCA) separately on the measures from each domain. PCA was performed on behavioral data from *n*=1537 training subjects (those without useable fMRI data but with no missing behavioral data). This transformation was then applied to the set of *n=*363 participants who had usable fMRI data, creating a PC score in each of the five domains for each participant.

#### fMRI data acquisition

Data analyzed were collected at one of two sites: Rutgers University Brain Imaging Center, or the Citibank Biomedical Imaging Center. Rutgers data were collected on a Siemens 3T Tim Trio magnet. Citibank Biomedical Imaging Center data were collected on a Siemens 3T Prisma. Both sites used the following parameters for the functional *Despicable Me* scan: TR=800ms, TE=30ms, # slices=60, flip angle=31°, # volumes=750, voxel size=2.4mm^3^, multiband 6. For more information regarding Healthy Brain Network scan parameters, please see (Alexander et al., 2017) and http://fcon_1000.projects.nitrc.org/indi/cmi_healthy_brain_network.

#### fMRI Preprocessing

AFNI was used to preprocess fMRI data. First, three volumes were removed from each run, followed by despiking and head motion correction. Then, functional images were aligned to the skull-stripped anatomical image with a linear transformation and then to the MNI atlas via nonlinear warping. Covariates of no interest were regressed from the data, including a 24-parameter head motion model (6 motion parameters, 6 temporal derivatives, and their squares) and mean signal from subject-specific eroded white matter and ventricle masks and the whole brain. Finally, images were band-pass filtered from .01 to .1 Hz.

### Dataset 2: Yale Attention

#### Participants

To compare our findings in development with that of an independent adult sample, we analyzed a dataset of healthy young adults who participated in a two-session neuroimaging experiment (Rosenberg et al., 2020; Yoo et al., 2022). Sessions were separated by 17.31 days on average (*s.d.* = 20.21 days, median = 12 days; Yoo et al., 2022). This project was approved by the Yale University Human Subjects Committee. Relevant to the present analysis, the participants in this study completed a fMRI run during each session in which they watched the short film *Inscapes* (Vanderwal et al., 2015). This film consists of dynamic visuals with no discernable narrative, making for an interesting comparison to the plot-driven clip shown in the Healthy Brain Network dataset. Prior to our access of the data, 33 participants were excluded due to unacceptable head motion (>3 mm maximum head displacement and > .15 mm mean framewise displacement), task performances falling 2.5 standard deviations above or below the mean, or because of low-quality imaging data. After downloading the data, an additional 9 scans were dropped due to having greater than 50% of frames censored for motion. This yielded a final sample of *n*=71 participants (47 F, 24 M; mean age = 22.86 ± 4.28 years, range = 18–36 years).

#### Behavioral data

Whereas the Healthy Brain Network behavioral data consists primarily of questionnaires, the Yale Attention dataset includes performance measures from three validated cognitive tasks. Consequently, we were able to determine if brain-behavior relationships differed across these different ways of assessing behavior. In this study, participants completed two sessions of the following three tasks designed to assess working memory and attentional performance: Gradual-onset continuous performance task (***gradCPT***), visual short-term memory task (***VSTM***), multiple object tracking task (***MOT***). For detailed descriptions of timing and other parameters for all tasks, see (Yoo et al., 2022).

The ***gradCPT*** (Esterman et al., 2013) assesses participants’ sustained attention and inhibitory control function. In this task, city and mountain photographs are displayed and participants were instructed to press a button in response to city scenes (appearing on 90% of trials) and withhold responses when mountain scenes appear (10% of trials). Images gradually transition from one to the next at a rate of 800 ms/trial. Performance was assessed by mean sensitivity (*d’*).

The ***VSTM*** task is a change detection task measuring visual working memory. During this task, participants viewed an array of 2, 3, 4, 6, or 8 colored circles, which were randomly positioned on the screen. After 100 ms, the circles were replaced by a fixation square for 900 ms before reappearing. In half of the trials, the colors of the circles remained unchanged, and on half the circles reappeared in a different color. Participants had to press one button if they detected a color change and a different button if there had been no change. Performance on this task was measured with Pashler’s K (Pashler, 1988).

Finally, the ***MOT*** assesses attentional selection and tracking (Luck & Vogel, 1997) and was adapted from code from Liverence & Scholl (2012). On each trial, 12 white circles appeared. Three or five of the circles (the targets) blinked green briefly before all circles began to move around the screen. Participants had to track the locations of the targets until the display stopped moving. When it stopped, one circle flashed, and participants pressed a button to indicate if that circle had been a target. Performance was assessed by mean accuracy across all trials.

#### fMRI data acquisition

Data were collected at the Yale Magnetic Resonance Research Center and Brain Imaging Center on a 3T Siemens Prisma with a 64-channel head coil. The following parameters were used for the functional scan analyzed here (*Inscapes* movie-watching): TR=1,000ms, TE=30ms, # slices=52, flip angle=62°, # volumes=600, voxel size=2.5mm^3^, multiband 4. For more information regarding the scan parameters of this data set, please see (Yoo et al., 2022).

#### fMRI Preprocessing

AFNI was used to preprocess fMRI data. First, three volumes were removed from each run. Then censoring was performed, removing volumes in which greater than 10% of voxels were outliers, or for which the Euclidean norm of the head motion parameter derivatives was greater than .2 mm. Despiking, slice-time correction, motion correction, and regression of mean signal from the CSF, white matter, and whole brain was performed. Additionally, 24 motion parameters (6 motion parameters, 6 temporal derivatives, and their squares) were regressed from the data. Functional images were aligned to the skull-stripped anatomical image with a linear transformation, and to the MNI atlas via nonlinear warping.

#### Pairwise Intersubject Time-course Correlation

For every individual in each dataset, preprocessed BOLD signal time-courses were averaged across voxels within regions of interest using a 268-node whole-brain parcellation (Shen et al., 2013). For every node, each participant’s BOLD signal time-course was *z*-scored within-subject and Pearson correlated with that of every other individual in the cohort. This yields a participant-by-participant “brain-similarity” matrix for each node, with each cell representing the synchronization of activation in that node for two individuals. We repeated this process in the Healthy Brain Network sample and the Yale Attention sample.

#### Intersubject Representational Similarity Analysis

After creating intersubject correlation matrices for each node and dataset cohort, we implemented intersubject representational similarity analysis (IS-RSA) to test two models of brain-behavior relationships: 1) a *Nearest Neighbors* model hypothesizing that individuals with similar behavioral PC scores will show greater brain similarity and 2) an *Anna Karenina (AnnaK)* model stating that individuals with high behavioral scores will show greater brain similarity whereas low scorers will show lower brain similarity (or vice versa) (Chen et al., 2020; Finn et al., 2020; van Baar et al., 2019). The goal of this approach is to determine which of the models, if either, explain the observed relationship between brain similarity and behavioral similarity.

To do this we computed three participant-by-participant matrices: one for brain similarity (pairwise intersubject correlation), and two for behavioral similarity (see Figure 1). For the *Nearest Neighbors* model, behavioral similarity is defined as the absolute value of the difference in behavioral scores. For the *AnnaK* model, behavioral similarity is the mean of the behavioral scores.

**Fig 1:**
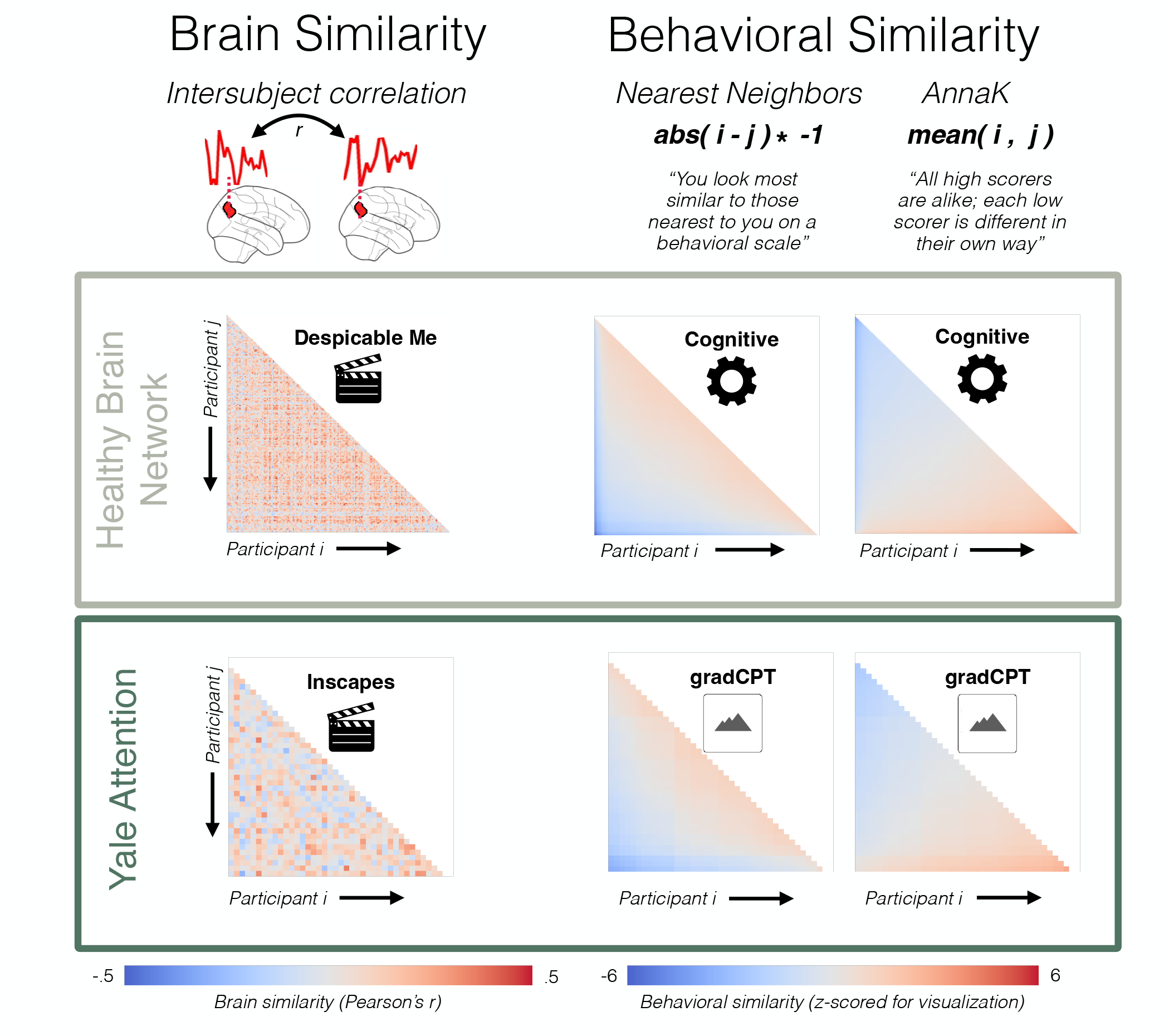
Brain and behavioral similarity matrices. Here we show similarity matrices, where each row and each column represent a participant. Participants are ordered by behavioral score (ascending). On the left are brain similarity matrices, defined as pairwise intersubject correlation for an example node (node 41) in the Shen parcellation for each dataset. On the right are two different behavioral similarity matrices for the first cohort of each dataset. The first behavioral matrix is calculated according to the *AnnaK* model, while the second is calculated according to the *Nearest Neighbors* model. To assess the relationship between brain similarity and behavioral similarity, we run intersubject representational similarity analysis Spearman correlating the brain and behavioral matrices.

To test how well our models capture brain-behavior relationships, we apply Spearman partial correlation to relate the vectorized lower triangles of the symmetric brain and behavioral similarity matrices, controlling for participant age and sex. To control for participant age and sex, we created three additional matrices: 1) a sex matrix (where a cell contains a one if participant *i* and *j* have the same sex, zero otherwise), 2) an *AnnaK* age matrix (where a cell contains the average of participant *i* and participant *j*’s ages), and 3) a *Nearest Neighbors* age matrix (where a cell contains the absolute value of the difference of participant *i* and participant *j*’s ages). These control matrices were regressed out of the brain and behavioral similarity matrices, and the subsequent analysis was performed on the residuals. Ultimately, the primary measure of interest is the Spearman correlation between the residualized brain and behavioral similarity matrices (calculated using a Python implementation of the Mantel test https://github.com/jwcarr/mantel). This correlation indicates how well the *AnnaK* and *Nearest Neighbors* models, respectively, fit the observed brain data. This correlation is performed in each of the 268 nodes in our parcellation for the IS-RSA of intersubject correlation.

To determine significance of model fit for each node, we performed permutation testing (10,000 permutations). For each permutation, the subject labels of the behavioral matrix were randomly shuffled, and the correlation between brain and behavioral matrices is re-calculated, generating a null distribution of 10,000 Spearman rho-values for each node. After using the null distribution to calculate a two-tailed *p*-value for every node, the Holm-Bonferroni method was used to correct for multiple comparisons. In order to assess the consistency of each model’s effects, we repeated the stated analyses in a split-half fashion. Participants were randomly divided into two cohorts, and intersubject RSA was performed separately on each cohort. This analysis allows us to determine how similar the rho-values for a given node are across the two cohorts.

To test whether there is greater evidence across the whole brain than would be expected by chance, we used an approach akin to the familywise error control method described in Finn et al., 2020. We randomly generated *p*-values for each of our 268 nodes, calculated the number of nodes which survive a *p*-threshold of .05, and repeated this 10,000 forming a null distribution. We compare the observed number of significant nodes to this null distribution to obtain a whole-brain *p*-value.

Finally, we directly compare *AnnaK* and *Nearest Neighbors*, testing if we find greater evidence for either model when averaging effects across the whole brain. To do so, we first took the difference in the Fisher-Z transformed rho-values for each node (subtracting the *Nearest Neighbors* Fisher-Z value from the *AnnaK* Fisher-Z value). Next, we averaged this difference across all 268 nodes to get a single difference value, at which point we applied the inverse Fisher-Z transform. To assess significance, we repeated this process 10,000 times, each time shuffling the subject labels of the behavioral matrix, to create a null distribution. Using this null distribution, we calculate a two-sided, non-parametric *p*-value for each behavioral task in each of our two datasets.

## Results

### Principal component analysis of behavioral measures in Healthy Brain Network data

After dividing the behavioral measures into five broad behavioral domains (see Supplementary figure 2), we applied principal component analysis (PCA) to the measures in each behavioral domain in the Healthy Brain Network sample (for cognitive task-specific analyses, see Supplementary Figure 3). We examined the resulting components, finding that the first principal component explained 47%, 39%, 40%, 68%, and 47% of variance, for the Cognitive, Social, Emotional, Attention, and Language domains, respectively (Figure 2). Since the first principal component explains a large portion of the variance, we retained this component as a summary score for each domain and used these summary scores in the remainder of our analyses.

**Fig 2:**
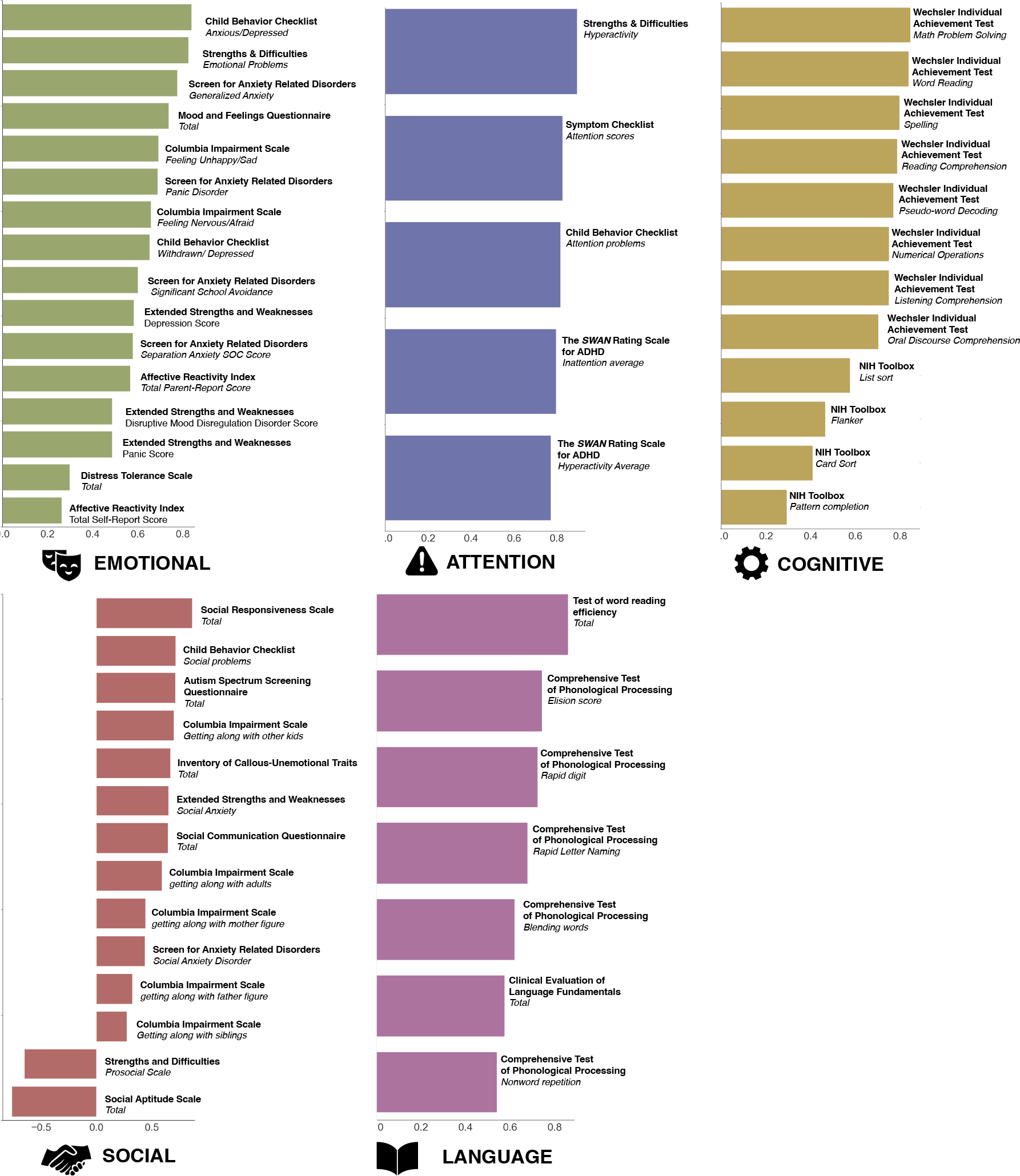
First principal component of five behavioral domains in the Healthy Brain Network sample. The contribution of each variable to the first principal component of each of the five domains of behavior analyzed in the Healthy Brain Network sample. In each plot, the x-axis shows the correlation between the original measure and the first principal component. In later analyses, Emotion, Attention, and Social scores are reverse coded for visualization, such that a greater value indicates better function in each respective domain.

Generally, the first principal component demonstrated a negative loading for the variables indicative of worse function (e.g., measures of depression symptoms in the Emotion domain) and a positive loading for variables indicative of better function. For the domains where the opposite was true (“negative” variables loaded positively, while “positive” variables loaded negatively), we multiplied the PC score by negative one. Consequently, in all domains a higher score indicates better function in that domain. Notably, the Cognitive and the Language components are the only domains that include behavioral task scores; social function, emotional function, attention contain only self-report or parent-report questionnaires. We did not apply PCA to the behavioral measures from the Yale Attention dataset as there were only three measures total, which assessed aspects of attention and working memory.

### Pairwise Intersubject Time-course Correlation

To ask if we can better understand individual differences in behavior by examining intersubject correlation (ISC), we first need to establish that the fMRI BOLD activity time courses of the movie watching scans are correlated, as we would anticipate. To perform this “sanity check,” we averaged ISC values across all pairs of participants for each node in our 268 node parcellation (Supplemental Figure 1). Average ISC across the whole brain was positive in both datasets and no nodes exhibited negative average ISC (Healthy Brain Network: mean *r*=.062, range: *r*=[.008, .249]; Yale Attention: mean *r*=.038, range: *r*=[.014, .268]). Overall, we observe the highest levels of ISC in regions predicted to be the most synchronized during movie watching: areas associated with visual and auditory processing. For instance, in the Healthy Brain Network sample, which included an audiovisual movie stimulus, the nodes with the five highest ISC values were in one of three areas: extrastriate cortex (*r*=.11, .13), Wernicke’s area (*r*=.10, .18), and visual association cortex (*r*=.12). In the Yale Attention sample, which included a visual-only movie, the five nodes with the highest ISC were all within primary visual cortex (*r*= .19, .20, .22, .25, .27).

### Intersubject representational similarity analysis

#### Intersubject RSA reveals significant representational similarity across behavioral domains

Previous work analyzing a working memory task in young adults demonstrates greater evidence for an *AnnaK* model than the *Nearest Neighbors* model (Finn et al., 2020). In other words, participants who scored similarly did not always display greater neural synchrony during movie watching. Rather, high scorers on the working memory task appeared more synchronized with high scorers and while low scorers were less in-sync with all others. Here we first ask whether this result is idiosyncratic to either the particular sample studied or to the particular type of movie analyzed. For instance, is the *AnnaK* effect dependent on the video shown containing a narrative arc? One possibility is that participants who score highly on a behavioral measure may interpret the narrative similarly in a way that is reflected in their BOLD activity. If this is the case, we would not expect to find evidence for the *AnnaK* model in the Yale Attention dataset, which uses *Inscapes*, a film with no narrative or plot to interpret. We next ask which model best explains the relationship between brain similarity and similarity in different types of behaviors. One possibility is that the *AnnaK* model will generally outperform the *Nearest Neighbors* model across a variety of behavioral domains. Alternatively, it is possible this effect is specific to working memory function, the measure assessed in previous work using data from the Human Connectome Project (Finn et al., 2020). If the *AnnaK* model is primarily a good fit for working memory measures, we would expect find the *AnnaK* model to be a good fit for only two out of the eight behavioral measures analyzed presently (the Healthy Brain Network cognitive domain and Yale Attention VSTM). Furthermore, since prior work used an adult sample, it is also possible that the *AnnaK* model does not describe brain–behavior relationships in development. In this case we would not expect to find an *AnnaK* effect in the Healthy Brain Network cognitive domain.

To investigate, we analyzed brain-behavior relationships in eight behavioral measures using two separate movies viewed in scanner and two independent samples (see Figure 3). In the Healthy Brain Network sample, the *AnnaK* model captures brain-behavior relationships in more nodes than would be expected by chance for both the Cognitive and Language domains (familywise *p <.*0001). Of these, relationships in five nodes in the Cognitive domain and one node in the Language domain nodes survive Bonferroni correction for 268 comparisons. Conversely, the *Nearest Neighbors* model does not capture brain-behavior relationships in more nodes than expected by chance for any domain (familywise *p* > .95), and no nodes show a significant *Nearest Neighbors* relationship after multiple comparisons correction. For four out of five behavioral domains, we observe more significant nodes (uncorrected *p*<.05) under the *AnnaK* model than the *Nearest Neighbors* model (difference in # of significant nodes, *AnnaK* – *Nearest Neighbors:* Cognitive: 135, Emotional: −2, Social: 11, Language: 27, Attention: 5). Finally, examining the split-half consistency of the effects for each model, the *AnnaK* model exhibits greater replicability than the *Nearest Neighbors* model for all five domains (*AnnaK* r_cohort 1, cohort 2_ Cognitive: .56, Emotional: .07, Social: .24, Language: .29, Attention: .23, *Nearest Neighbors* r_cohort 1, cohort 2_: Cognitive: 0, Emotional: -.1, Social: .01, Language: -.05, Attention: .01).

**Fig 3.**
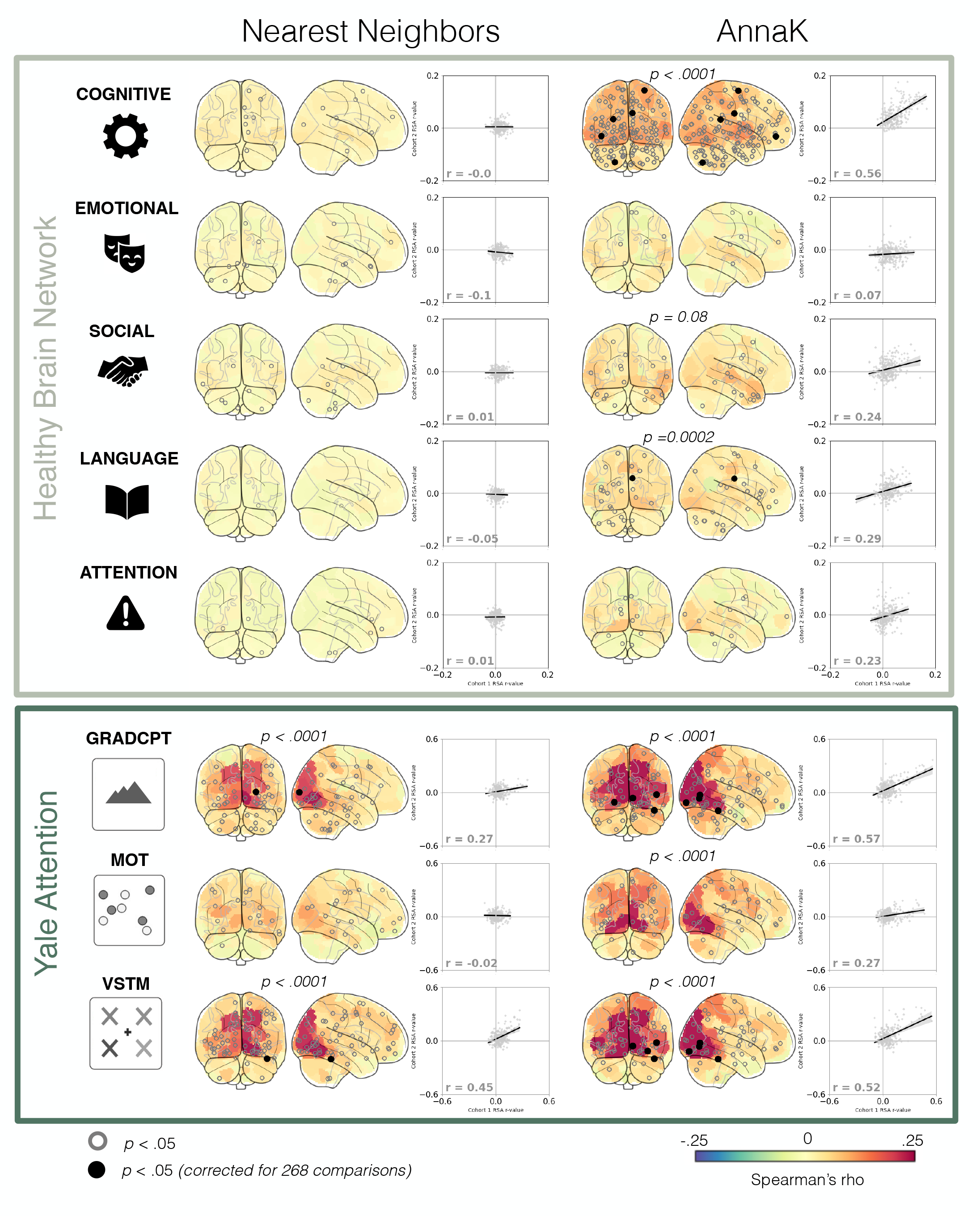
Relating behavior and neural synchrony with intersubject RSA. In the glass brain, nodes are colored according to representational similarity for each model in our two datasets. Nodes with a black circle (filled in) at their centroid demonstrate significant representational similarity (*p* <. 05, after correction for 268 comparisons). Nodes with a gray circle (empty) are significant before multiple comparisons correction (*p* < .05). *P*-values above the glass brains correspond to the whole-brain familywise p-value (see Methods). Glass brains without *p*-values above all show familywise *p* > .1. The scatterplots to the right of the glass brains show the reliability of the effects as determined by a split-half analysis. The *x*-axis is determined by the representational similarity rho-value in cohort 1 of the split-half analysis. The *y*-axis is determined by the representational similarity rho-value in cohort 2. The regression line and *r*-value labeled in the plot indicate the correlation of the effects across the two cohorts, providing insight into the consistency of the effects are across halves of the sample, with a higher correlation indicating greater consistency. Note: the rho-values in the scatterplot are separate from the Spearman’s rho node colors on the glass brain; the former represents effect consistency across all 268 nodes, while the latter denotes the fit of the model in that specific node.

In sum, we did not find support for *Nearest Neighbors*-style relationships in any domain. Across four out of five behavioral domains in the Healthy Brain Network sample, we found more support for the *AnnaK* model than the *Nearest Neighbors* model, and we found significant evidence for the AnnaK model in both the Cognitive and Language domains. This suggests that participants who score higher on assessments of cognitive function and language ability are more synchronized, in terms of brain activity, during movie watching while those who score lower are less synchronized.

The *AnnaK* model captures brain-behavior relationships in more nodes than would be expected by chance for both the Cognitive and Language domains (familywise *p* < .0001). More research is needed to determine if the result can be attributed to the use of task data in the Cognitive and Language domains (instead of self-report questionnaire data), or if it is due to the nature of the behavioral processes underlying the two measures. For instance, prior research examining a non-self-report social measure, number of social connections reported by one’s peers, found a robust *AnnaK*-effect in several brain regions (Baek et al., 2022).

Turning to the results in the Yale Attention Dataset, the *AnnaK* model reflected brain-behavior relationships in all three tasks, as evidenced by more significant nodes than would be expected due to chance (familywise *p <.*0001, see Figure 3). The *Nearest Neighbors* model significantly reflected brain-behavior relationships for gradCPT and VSTM (familywise *p <.*0001), but not MOT (familywise *p*= .13), performance. For two out of three tasks, we observe more significant nodes (uncorrected *p* < .05) under the *AnnaK* model than the *Nearest Neighbors* model (difference in # of significant nodes by domain, *AnnaK* - *Nearest Neighbors:* gradCPT: 4, MOT: 3, VSTM: −2). Of the two domains with nodes surviving comparisons for multiple corrections, both showed more significant nodes under the *AnnaK* model (difference in # of significant nodes by domain, *AnnaK* - *Nearest Neighbors:* gradCPT: 4, VSTM: 3). Finally, the effects of the *AnnaK* model demonstrated greater split-half consistency for all three behavioral tasks (difference in consistency by domain, *AnnaK* r_cohort 1, cohort 2_ - *Nearest Neighbors* r_cohort 1, cohort 2_: gradCPT: .30, MOT: .29, VSTM: .07). As in the Healthy Brain Network sample, results in the Yale Attention sample demonstrate greater evidence for the *AnnaK* model overall, wherein high scorers show greater similarity compared to high scorers and low scorers look dissimilar from everyone else.

Of all behaviors and models tested, the *AnnaK* model significantly reflects brain-behavior relationships for the most nodes in the Cognitive domain. There are more nodes demonstrating an *AnnaK*-effect for the Cognitive domain (143) than for any of the three cognition-related measures in the Yale Attention sample (gradCPT: 59, VSTM: 51, MOT: 31). One key difference between these measures is that the Cognitive measure incorporates data from several behavioral tasks (e.g., Flanker, List Sort) while the Yale Attention measures include data from just one behavioral task each. To ensure the robust *AnnaK* effect in the Cognitive measure cannot be attributed to summarizing across tasks, we took each variable comprising the Cognitive measure, and replicated the IS-RSA with these variables individually (Supplementary Figure 3). Out of the 12 variables tested, eleven demonstrate a better fit with the *AnnaK* model judging by the number of significant nodes. Consequently, the *AnnaK*-effect in the Cognitive domain does not appear to be an artifact of the choice to collapse across tasks.

As a final method of assessing the models, we computed a whole-brain summary score by averaging all nodes’ rho-values. Judging by the difference in this summary score, *AnnaK* model numerically outperformed the *Nearest Neighbors* model in every behavioral domain tested across both datasets. Comparing this difference score to a permuted null distribution of difference scores, the *AnnaK* model better captured the relationship between brain and behavioral similarity than the *Nearest Neighbors* model in the Cognitive domain in the Healthy Brain Network and in the gradCPT in the Yale Attention dataset (*p*=.002 and *p*=.022, respectively). Ultimately, relative to the *AnnaK* model, we found less compelling evidence for the *Nearest Neighbors* model, a model grounded in the intuitive hypothesis that people who are more similar on a behavioral scale will show more similar brain responses. Rather, on the whole, we found more evidence that brain-phenotype relationships examined showed an *AnnaK*-style pattern, wherein high scorers were synchronized with other scorers, while low-scoring participants showed less synchrony across the board. Although more research is necessary to determine under what conditions the *AnnaK* model describes brain-behavior relationships, here we demonstrate that this effect is robust to sample demographics (e.g., adults vs. adolescents), movie type (e.g., narrative vs. non-narrative), and the behavior itself (e.g., sustained attention vs. working memory).

### Anatomical distribution of intersubject RSA effects differs by dataset

Aside from how well the *AnnaK* and *Nearest Neighbors* models describe the relationships between brain and behavior, we were also curious as to *where* in the brain these models fit the data well. One possibility is that we find more significant nodes in areas where ISC is high; this would mean auditory processing areas for the Healthy Brain Network dataset and visual processing areas for both datasets (the Yale Attention stimulus did not have audio). Alternatively, if ISC is *too* high, it may not provide enough variance to successfully relate to individual differences in behavior. The nature of the stimulus type may also modulate where each model of representational similarity fits. For instance, synchrony in higher-order regions may prove more important when the video stimulus contains more emotional or narrative content, such as the clip shown to the Healthy Brain Network participants. Finally, the distribution of nodes could depend on the model in question, with some regions exhibiting more *AnnaK* or *Nearest Neighbors*-style relationships respectively.

To answer this question, we plot nodes demonstrating significant representational similarity (uncorrected *p* < .05) within a given model according to which network they belong to (Supplementary Figure 4). In the Cognitive domain, looking at the *AnnaK* model, the significant nodes are fairly evenly distributed across all networks. No one network contains more significant nodes than would be expected by chance, although the visual association network shows the greatest number of nodes. Alternatively, in the Yale Attention Sample, every behavior for both models shows the greatest number of nodes in early visual cortex. The number of significant nodes in primary visual cortex are greater than would be expected by chance for both the gradCPT and VSTM tasks across both models (all *p* < .001) and for the MOT task in the *AnnaK* model (*p* < .01). This pattern of results is consistent with the hypothesis that the narrative content of a video modulates where in the brain neural synchrony relates to individual differences, with significant nodes clustered in early visual cortex for the non-narrative but not for the narrative stimulus. However, a number of alternative factors may drive this difference between datasets, and as a result, more research is needed to conclusively parse how video content affects intersubject RSA.

### Model fit highly is correlated with intersubject correlation

To better understand how our power for detecting individual differences varies as a function of ISC, we also assessed to what extent the intersubject RSA model fit is correlated with average ISC (Supplementary Figure 5). If neural synchrony is associated with greater power for picking up on individual differences, we would expect a positive correlation between ISC and model fits. Indeed, this is what we observe. In every instance where the intersubject RSA was significant at the whole-brain level, we see a strong positive correlation between node-wise ISC and node-wise IS-RSA rho (*AnnaK*: Cognitive: r=.77 Language: r=.37, gradCPT: r=.87, MOT: r=.77, VSTM: r=.87, *Nearest Neighbors*: gradCPT: r=.77, VSTM: r=.80). In fact, in 13 out of 16 instances (2 models x 8 behavioral measures) we observe a positive correlation between ISC and IS-RSA effects. Here, perhaps counterintuitively, greater similarity across subjects’ brain activity appears to better allow us to understand behavioral differences. This is consistent with prior work showing that movie data is better than rest data for predictive modeling of individual differences in behavior (Finn & Bandettini, 2021). Movie watching may influence brain activity in a way that makes participants appear more similar, but it may also increase signal to noise. Future work remains to determine if there is an upper bound to this effect, such that too much neural synchrony results in a loss of power for modeling behavioral variability.

### Both AnnaK and Nearest Neighbors models significantly describe relationship between neural similarity and age in development

The *AnnaK* model significantly captured brain-behavior relationships in the Healthy Brain Network Cognitive and Language domains and for all three Yale Attention sample tasks. The *Nearest Neighbors* model significantly fit the data for the Yale Attention gradCPT and VSTM task. We next wanted to ask about the relationship between a different type of phenotypic measure—age—and neural synchrony. Do participants more similar in age show higher ISC as predicted by the *Nearest Neighbors* model, or do older (or younger) participants show more similar ISC as predicted by the *AnnaK* model? One might predict similarity in participant age to exhibit a *Nearest Neighbors*-style relationship, particularly in the Healthy Brain Network sample, due to its inclusion of ages that span the course of adolescence (6-21). For instance, a 7-year-old and a 21-year-old are likely to have different interpretations of a narrative stimulus, and these interpretations may be reflected in different fMRI time-courses during movie-watching. Unlike the developmental Healthy Brain Network sample, the Yale Attention dataset is a sample of young adults (aged 18-36). The brain undergoes significant changes during development, and shows relative stability during early adulthood (Bethlehem et al., 2022), and consequently we might anticipate less age-related variability in video processing in the Yale sample, potentially resulting in little age-related differences in neural synchrony. This in turn would result in a poor fit for both the *AnnaK* and *Nearest Neighbors* models.

Examining the results of the age-specific intersubject RSA (Figure 4), we find significant evidence of the *Nearest Neighbors* model in the Healthy Brain Network sample (241 significant nodes, 139 surviving multiple comparisons correction, familywise *p* < .0001). In line with our hypothesis, this implies that there is greater neural synchrony among participants that are closer in age. As for the *AnnaK* model, we observe, in contrast to our behavioral findings, primarily *negative* correlations between brain similarity and *AnnaK*-defined behavioral similarity (178 significant nodes, 17 surviving multiple comparisons correction, familywise *p* < .0001). This negative correlation implies that the older participants get, the more *dissimilar* they appear from each other, in terms of brain synchrony. In the Yale Attention dataset, by contrast, neither the *AnnaK* nor the *Nearest Neighbors* models well-described the relationship between brain and age similarity (13 and 14 significant nodes respectively, 0 surviving multiple comparisons correction, familywise *p*-values > .47).

**Fig 4.**
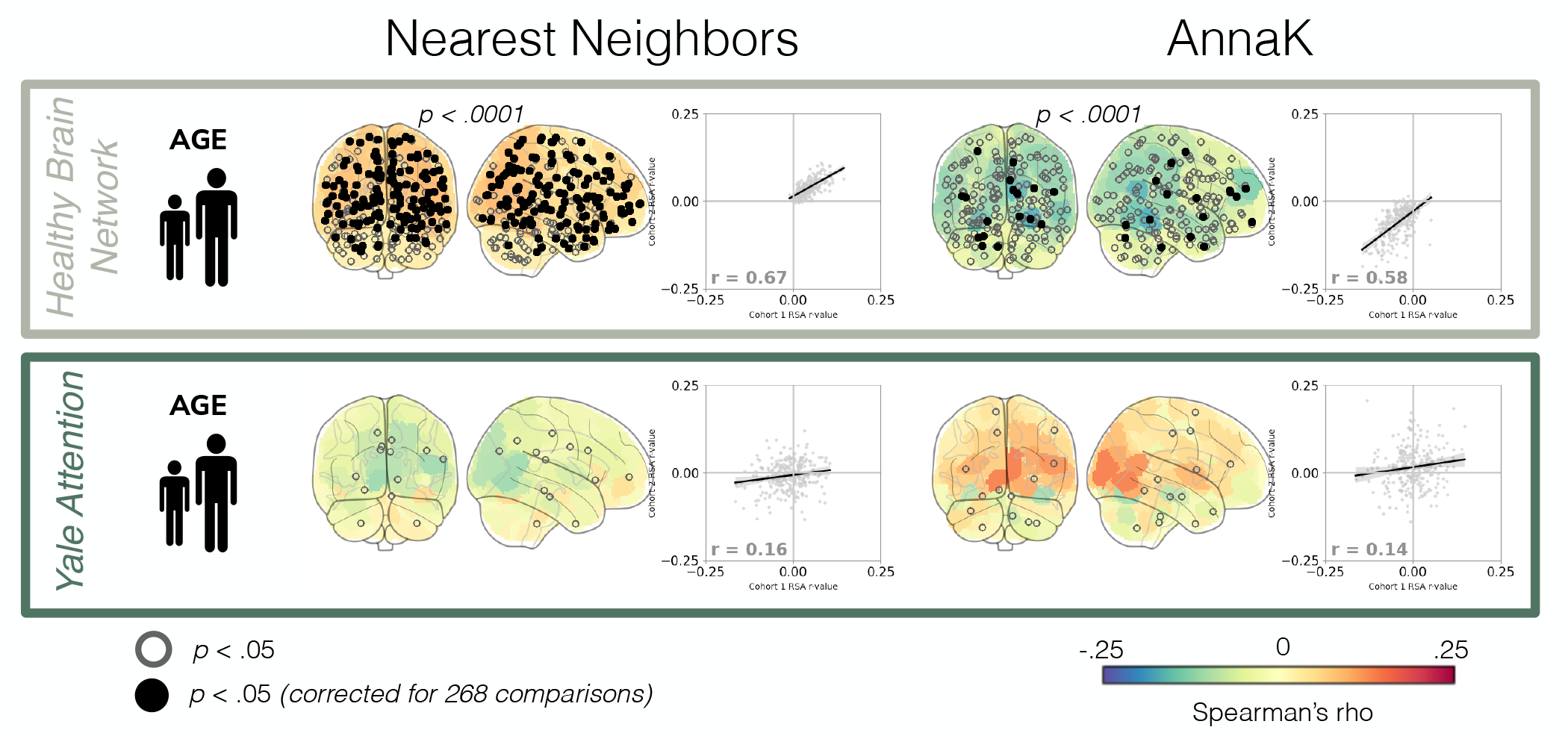
Relating age and neural synchrony with intersubject RSA. In the glass brain, nodes are colored according to representational similarity for each model in each dataset. Nodes with a black circle (filled in) at their centroid demonstrate significant representational similarity (*p* < .05, after correction for 268 comparisons). Nodes with a gray circle (empty) are significant before multiple comparisons correction (*p* < .05). *P*-values above the glass brains correspond to the whole-brain familywise *p*-value (see Methods). Glass brains without *p*-values above all show familywise *p* > .1. The scatterplots to the right of the glass brains show the reliability of the effects as determined by a split-half analysis.

In sum, both the *AnnaK* model and the *Nearest Neighbors* model describe the relationship between age and intersubject correlation in the Healthy Brain Network sample. This raises the possibility, initially proposed by Finn and colleagues (Finn et al., 2020), that for some phenotypic measures, modeling phenotypic similarity as a combination of the *AnnaK* and *Nearest Neighbors* models might yield the best fit to brain similarity (for instance, by using the formula *abs(i-j)* mean(i,j)*). In the Yale Attention dataset, neither model was a good fit, although more research is necessary to determine if this is due to the constrained age range of this sample or some other factor.

## Discussion

Recent work hypothesizes that similarities in behavioral traits are reflected in similar BOLD responses to movies, but that this relationship does not always hold constant across the behavioral spectrum (Finn et al., 2020). Specifically, Finn and colleagues found that participants who scored highly on a test of working memory showed greater neural synchrony with other high scorers, while individuals who scored low appear less synchronized across the board. In this instance, neural similarity is indicative of behavioral similarity, but only in high-scoring participants. How does the relationship between neural similarity and behavior play out in other behavioral domains?

In the present study, we investigate this question, examining brain-behavior relationships in eight behaviors across samples from two datasets: the Healthy Brain Network dataset (Cognitive, Attention, Social, Emotional, and Language) and the Yale Attention dataset (gradCPT, MOT, VSTM). Using intersubject-RSA, we tested two models of neural synchrony: 1. the *Nearest Neighbors* model, stating that individuals will look similar to those nearby on some behavioral scale and 2. the *AnnaK* model, stating that high scorers will look like other high scorers while low scorers will show greater variability (or vice versa). Overall, our analyses revealed greater evidence for the *AnnaK* model of brain-behavior relationships, particularly for the Cognitive domain in the Healthy Brain Network Dataset and the sustained attention task in the Yale Attention dataset. Put another way, results here suggest that in tasks across datasets and behavioral domains, high scorers tend to resemble high scorers, whereas low scorers appear dissimilar from everyone else. While the *Nearest Neighbors* model also fit the data above chance for two of the eight behavioral domains tested (gradCPT and VSTM), in no behavioral domain did it clearly outperform the *AnnaK* model when examining number of significant nodes and family-wise *p*-values. This finding is striking considering that the *Nearest Neighbors* model is arguably a more common way of operationalizing the association between brain activity and cognitive performance. Our results here replicate previous observations of *AnnaK*-style ISC-behavior associations (Finn et al., 2020) and underscore the importance of considering the meta-relationship between brain and behavior to avoid missing the forest for the trees.

Our findings here, as well as the results of Finn et al., 2020, provide substantial evidence of an *AnnaK*-style relationship in cognition-related domains. Intriguingly, however, the advantage of the *AnnaK* model is more apparent in the Healthy Brain Network and Human Connectome Project samples, and less clear in the Yale Attention sample.

One factor that that may drive this difference across datasets is the relative granularity of the behavioral measure. Perhaps it is not surprising that a “coarse” measure of behavior, such as the Cognitive PC score defined here, shows greater variability among low scorers. Whereas the highest scorers may perform relatively well on all tasks, the low-scoring group could be made up of individuals who performed poorly on different tasks, such as spelling and working memory. In this case, the variability among low scorers could be attributed to difficulties with spelling and working memory, respectively. To control for this potential explanation, we conducted IS-RSA separately on each component variable of the Cognitive PC score. Even when examining tasks individually, however, we continue to find greater evidence for the *AnnaK* model (Supplementary Figure 2). An additional strike against a “granularity” account is the robust evidence for the *AnnaK* model in the Human Connectome Project, where a single task, the NIH List Sorting working memory task, was analyzed by Finn et al. (2020). Another possibility is that the difference across datasets arises from the use of movie data from two fMRI sessions in the Yale Attention analyses, compared to data from one session in all other analyses. To investigate this, we replicated our IS-RSA analyses separately by session (Supplementary Figure 7). While effects are weaker in the second session, possibly due to lower ISC, we see largely similar patterns across sessions, making it unlikely that the multisession analysis explains the cross-dataset differences.

A third factor that may contribute to this discrepancy is the type of movie stimulus used. Human Connectome Project and Healthy Brain Network participants viewed video stimuli that included narratives, while the Yale Attention sample viewed an abstract film lacking narrative or representational content. Perhaps the variance in brain activity evidenced in those who scored lower on behavioral measures is heightened by the cognitive processes involved in processing narratives. Finally, one last factor that may influence differences in how the *AnnaK* model captures behavioral similarity across datasets is sample heterogeneity. In addition, both the Human Connectome Project and Healthy Brain Network include data from multiple scanning sites and, in the case of the Healthy Brain Network, intentionally focus recruitment on participants with symptoms of psychopathology. The Yale Attention dataset, alternatively, includes a smaller non-clinical sample of adult participants. If sample heterogeneity proves to be an important factor to brain-behavior similarity, as fMRI datasets grow larger and span a more diverse range participants, it may become increasingly difficult to understand behavior through analyses which assume a one-to-one mapping between brain and behavioral similarity.

When relating intersubject-RSA to age, we observed evidence for the *Nearest Neighbors* and *AnnaK* models in the Healthy Brain Network sample only. These results suggest that, in the Healthy Brain Network dataset, younger participants and participants closer in age show greater neural synchrony. While this result may seem curious when juxtaposed with our behavioral findings, it is in line with prior work demonstrating that ISC as measured with EEG decreases with age (participants aged 5-44) (Petroni et al., 2018) and ISC as measured with fMRI decreases with age (participants aged 18–88) (Campbell et al., 2015). However, existing literature is somewhat mixed with regard to the association between neural synchrony and age, with some evidence suggesting that adults exhibit greater ISC compared to children, particularly in the default mode network (Moraczewski et al., 2020). Another study analyzing data from the Healthy Brain Network found, in line with our findings here, more areas of the brain where ISC was greater in younger participants than older participants (Cohen et al., 2022). Inconsistent with the current findings, however, the same study reports higher ISC among older participants in the auditory cortex (Cohen et al., 2022), and another which reports greater synchrony among older participants across the cortex more generally (Camacho et al., 2023). One factor that may contribute to this discrepancy is our use of a relatively conservative motion threshold (mean framewise displacement < .15 mm) in determining our sample. While this criterion should help safeguard against spurious motion-driven effects, we cannot rule out the possibility that motion in this sample is related to another variable affecting ISC (Power et al., 2012). Additionally, our analyses include all ages available in the Healthy Brain Network dataset (ages 6-21 years); when excluding participants 16 years and older (as in Camacho et al., 2023), we observe a primarily positive relationship between age and intersubject correlation (see Supplementary Figure 8). Consequently, future work remains to elucidate how ISC changes across development.

The *AnnaK* model may help us to characterize the nature of brain-behavior similarity, but future research is needed to determine the source of increased variance for one side of a behavioral scale. How can we understand features characteristic of the end of the spectrum marked by greater heterogeneity? One approach which may prove useful is the classification of behavioral performance. Among individuals with the same summary score on a behavioral task, there may be clusters of participants that can be differentiated based on their patterns of responses. For instance, perhaps some individuals take longer to recover after making errors. A separate group of participants may quickly regain focus after errors but show a marked deterioration in performance near the end of the task, as their concentration wanes. If neural similarity is higher within these clusters, but lower across them, this could help explain why there is more variation in brain activity among low-scoring individuals. However, there may be a limit to how well we can characterize this variability using individual differences approaches, if low-scoring individuals appear not only dissimilar to each other, but dissimilar to themselves. This may prove to be the case, considering recent work showing that intra-individual variability is associated with worse task performance, both in terms of session-to-session whole-brain functional connectivity (Corriveau et al., 2022) as well as trial-by-trial patterns of hippocampal activity (Poh et al., 2022).

The present study aims to further elucidate the connection between similarity in brain activity and similarity in behavior. Using intersubject representational similarity analysis, we conceptually replicate prior work, and empirically assess assumptions in cognitive neuroscience research which may otherwise go untested. In continuing to map the space of neural and behavioral similarity, future work will reveal the structure governing how neural activity produces our idiosyncrasies as individuals.

## Acknowledgments.

This work was supported by National Science Foundation BCS-2043740 (M.D.R.) and resources provided by the University of Chicago Research Computing Center. Data collection for the Yale Attention Dataset was supported by National Institutes of Health MH 108591 and National Science Foundation BCS 1558497.

## Data and Code Availability

Healthy Brain Network data are available at http://fcon_1000.projects.nitrc.org/indi/cmi_healthy_brain_network/. Yale Attention data are available at https://nda.nih.gov/edit_collection.html?id=2402. Code will be made available on GitHub upon publication at https://github.com/tchamberlain/nn_annak.

## Supplementary Information

**S1:**
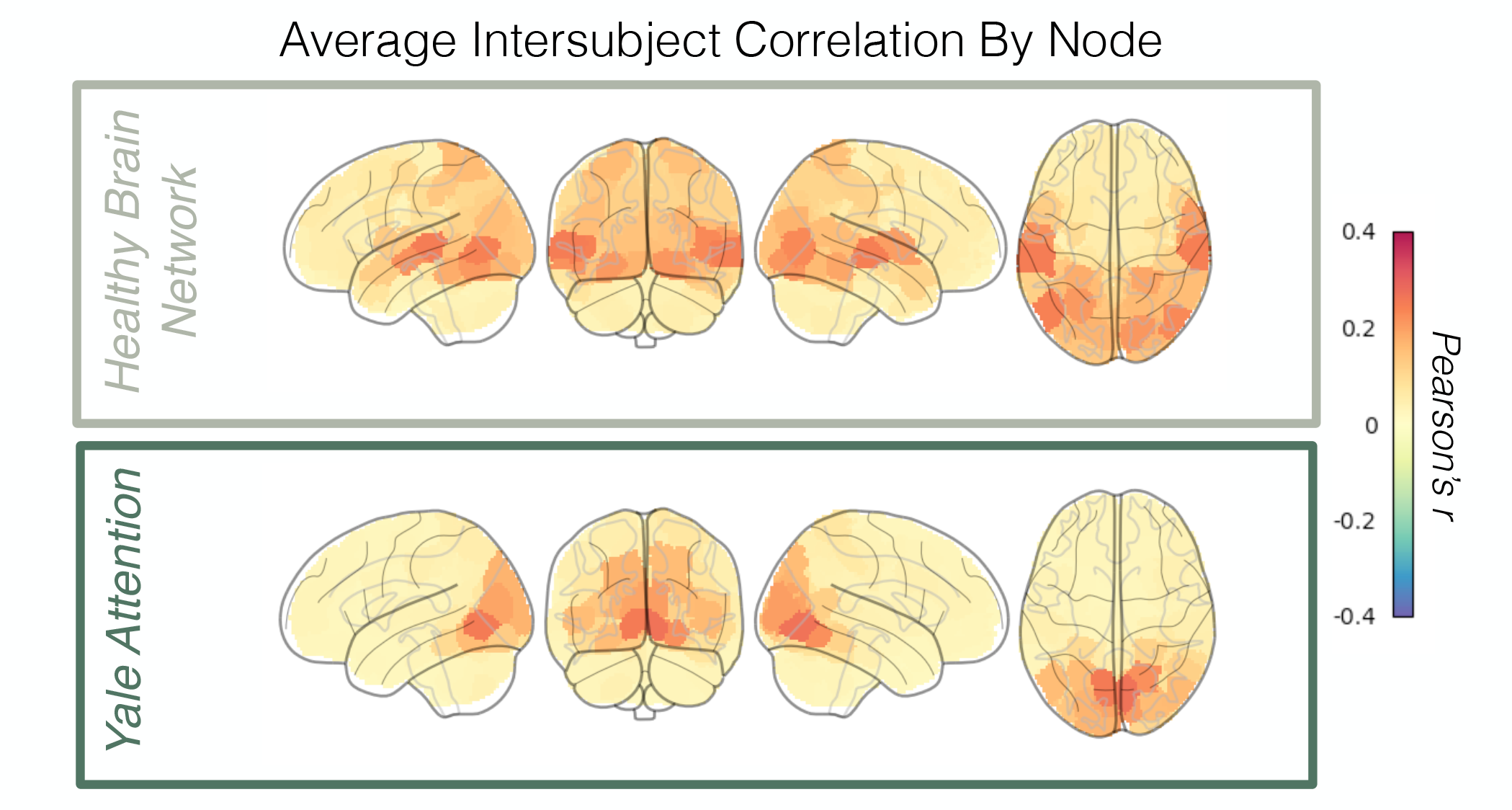
Intersubject correlation. Here we show pairwise intersubject correlation, calculated separately in each node in our 268-node parcellation, and averaged across all participants. Intersubject correlation is defined as the Pearson correlation between the BOLD time series of two different participants for a given node.

**S2:**
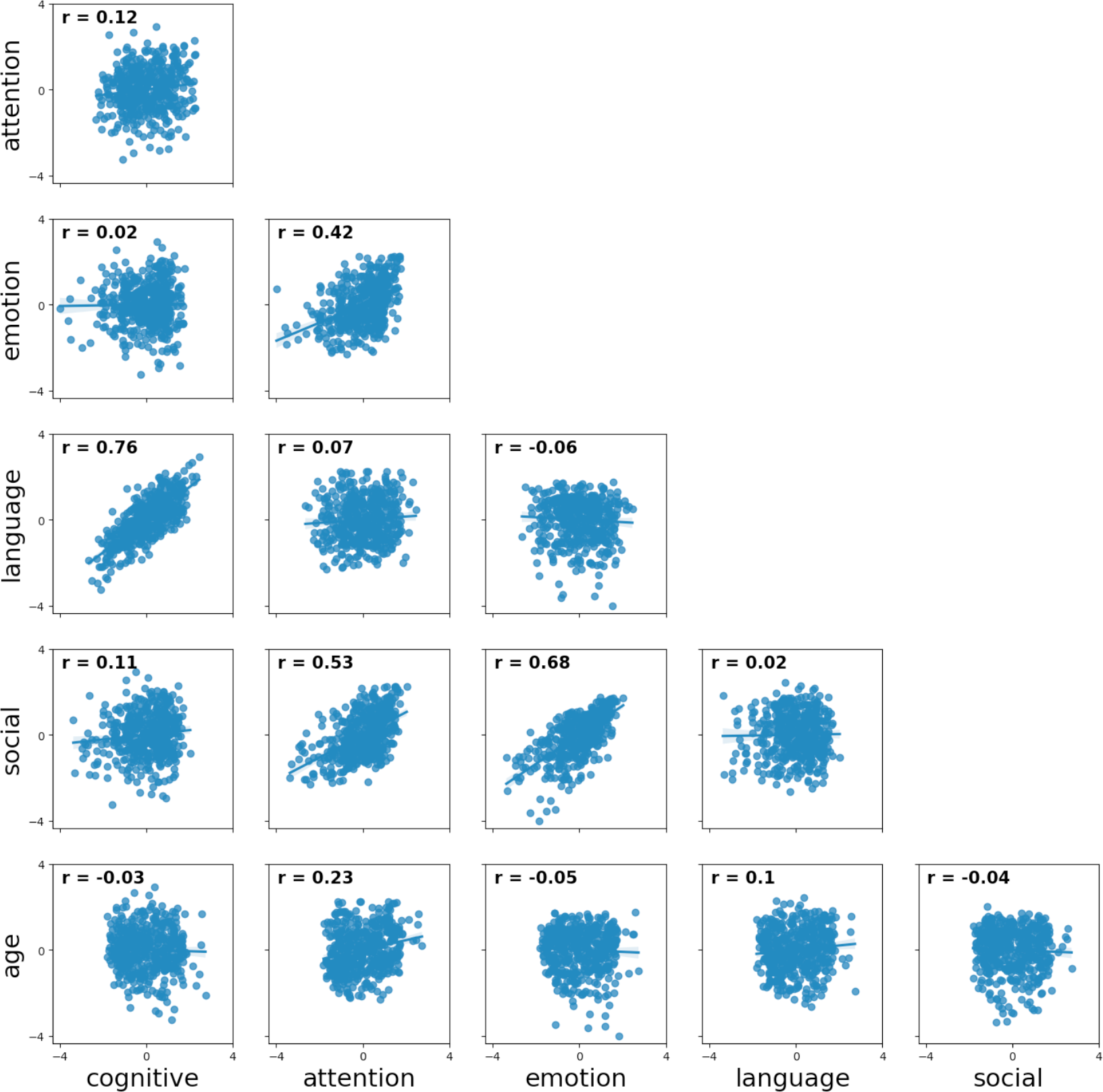
Associations between Healthy Brain Network behavioral scores. Scatterplots showing the relationship between z-scored behavioral scores in the Healthy Brain Network biobank. The r-value is the Pearson correlation between the two behaviors.

**S3:**
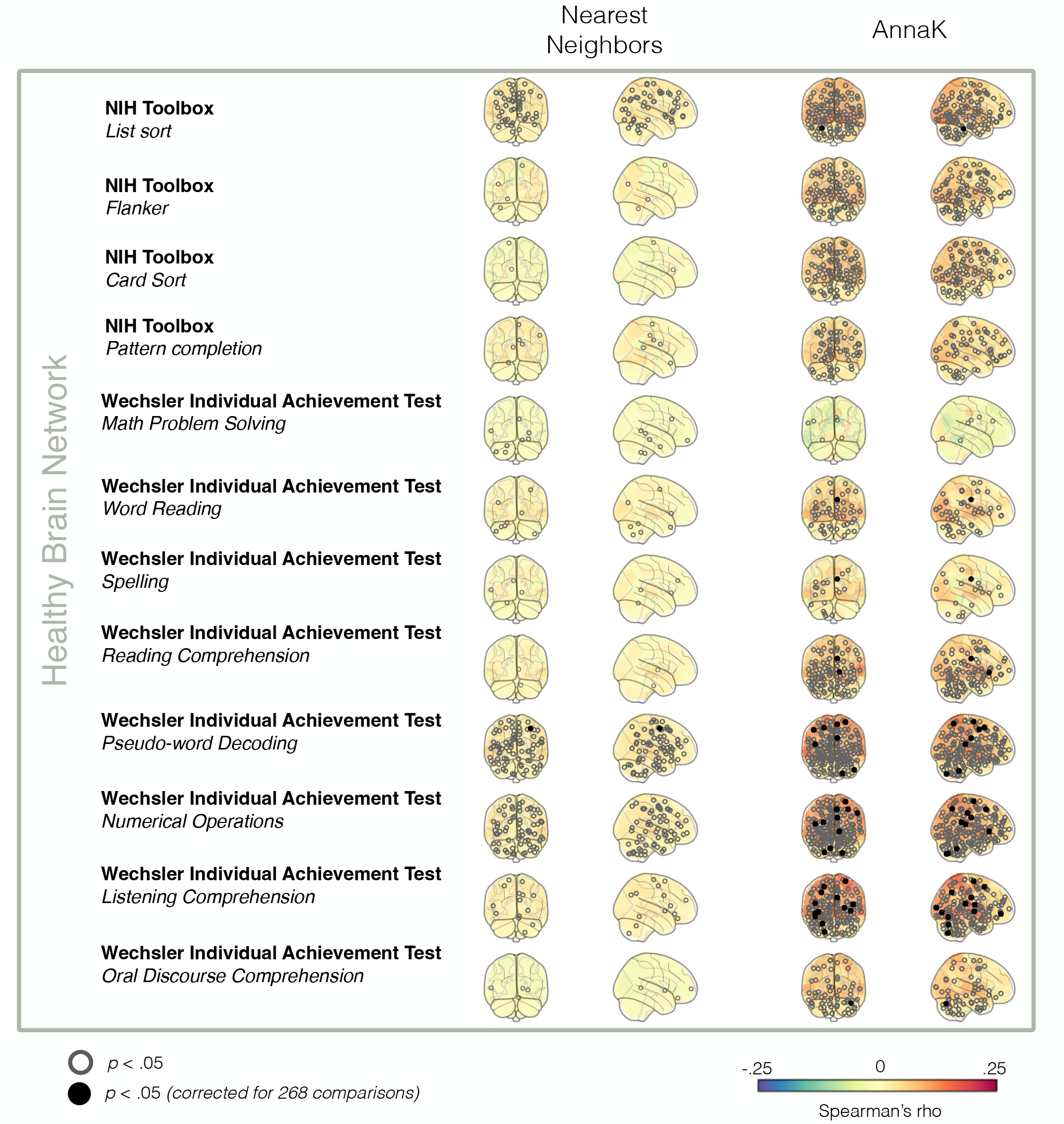
Intersubject RSA (Healthy Brain Network cognitive measures). IS-RSA results for each variable in the “cognitive domain” PCA. Plotting conventions are the same as figures in the main text (see Figure 3).

**S4:**
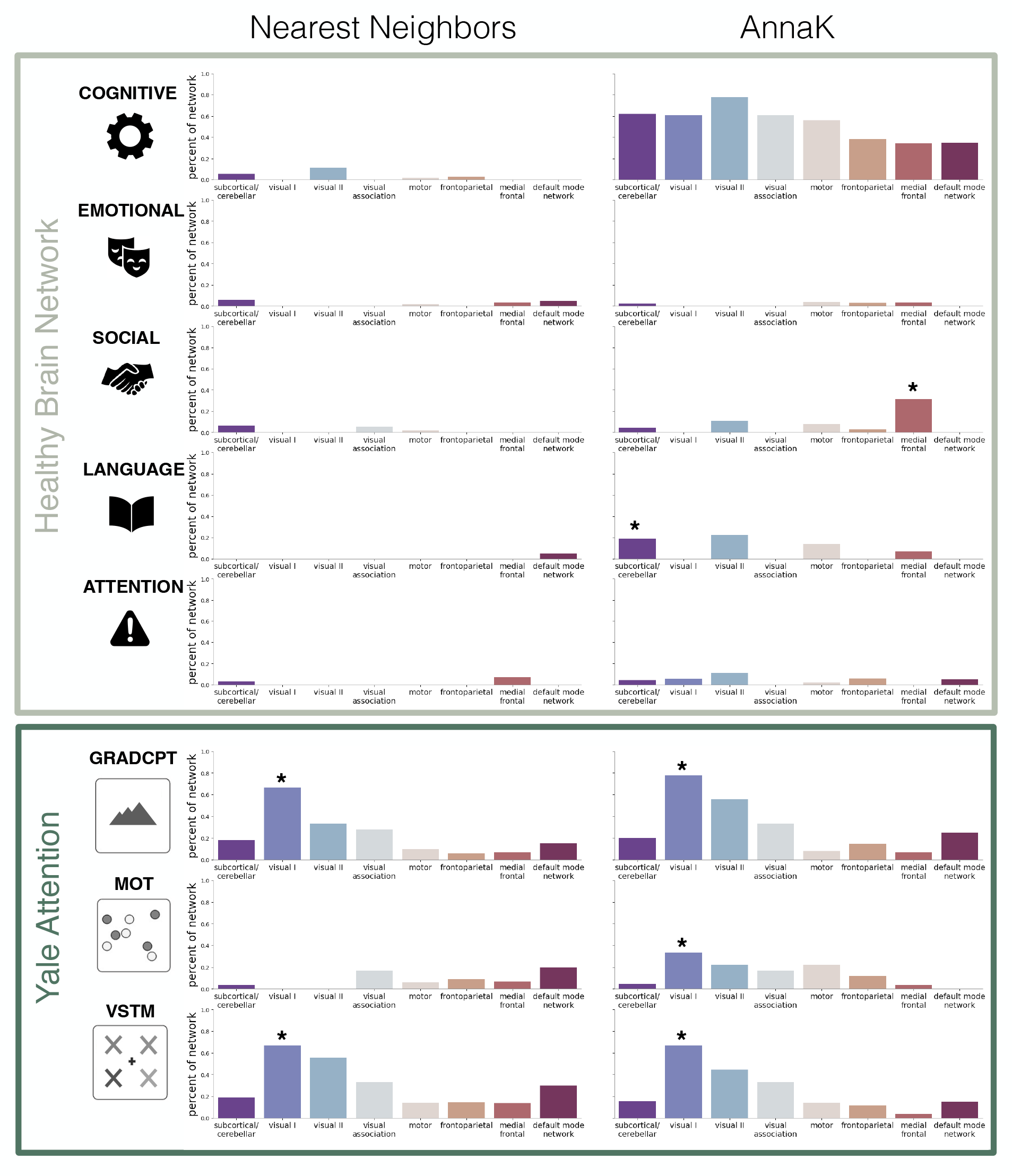
Intersubject RSA results by network. For each behavioral task, for each model, we show the percent of nodes in a given network that show significant representational similarity. Stars indicate that the given network demonstrates a greater number of nodes than expected by chance, as determined by permutation testing (p<.05 corrected for 8 comparisons).

**S5:**
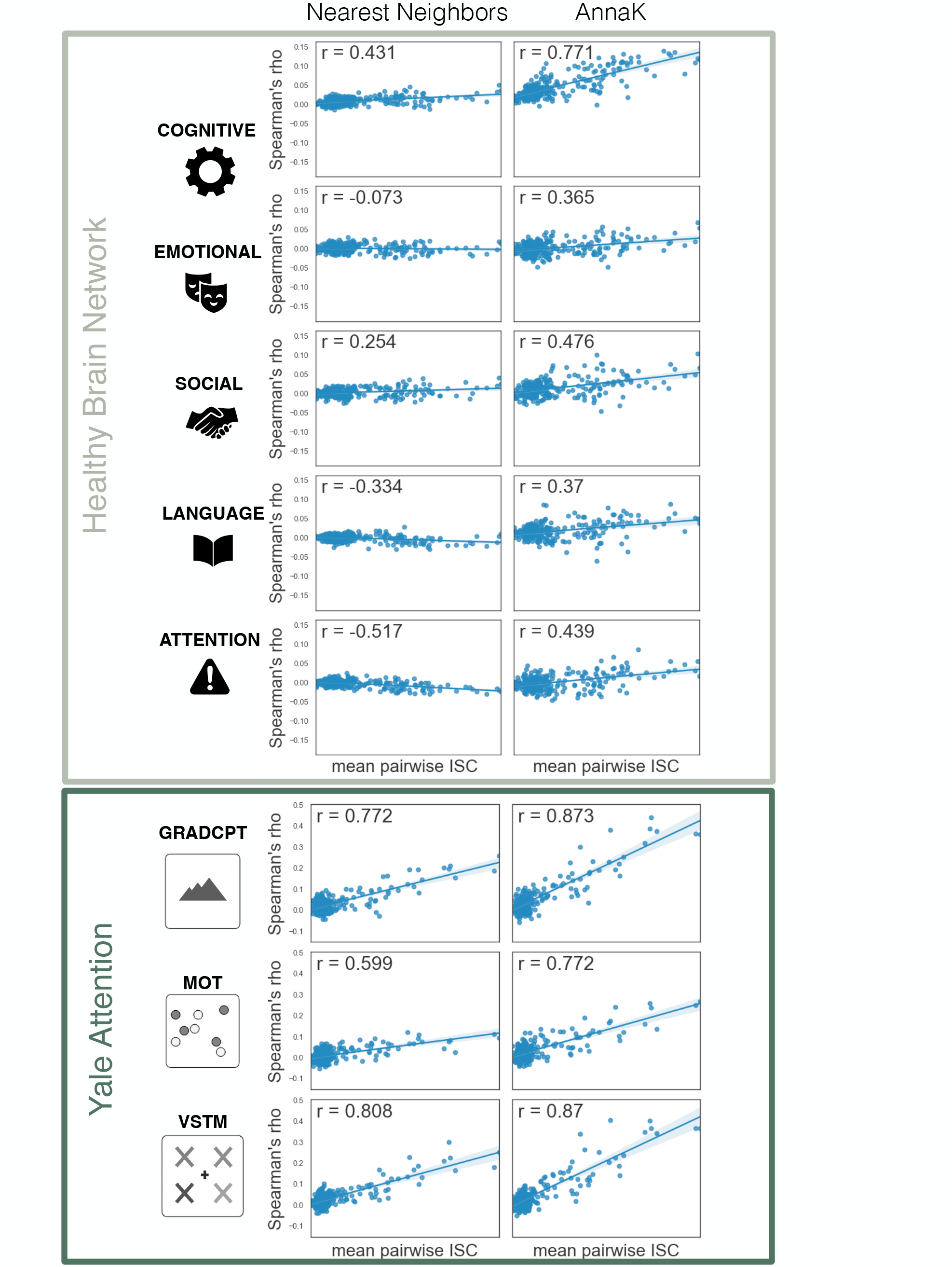
Correlation between IS-RSA Spearman R and ISC by node. Each point on each scatterplot is one of 268 nodes in the whole brain parcellation. The y-axis shows the r-value representing model fit in that node (brain similarity correlated with behavioral similarity for a given model). The x-axis shows the mean pairwise intersubject correlation in the node.

**S6:**
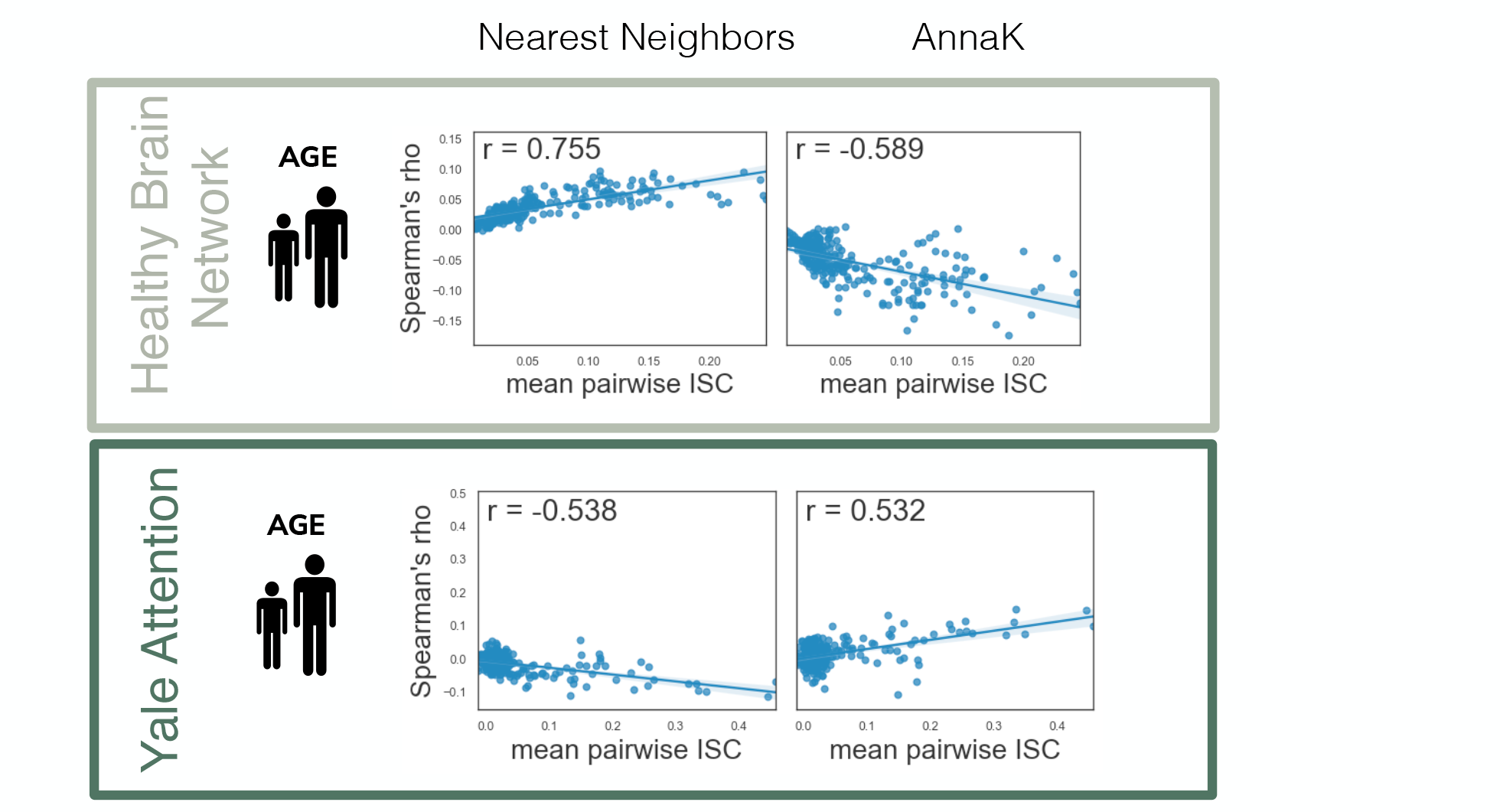
Correlation between IS-RSA Spearman R and ISC by node. Each point on each scatterplot is one of 268 nodes in the whole brain parcellation. The y-axis shows the *r*-value representing model fit in that node (brain similarity correlated with behavioral similarity for a given model). The x-axis shows the mean pairwise intersubject correlation in the node.

**S7:**
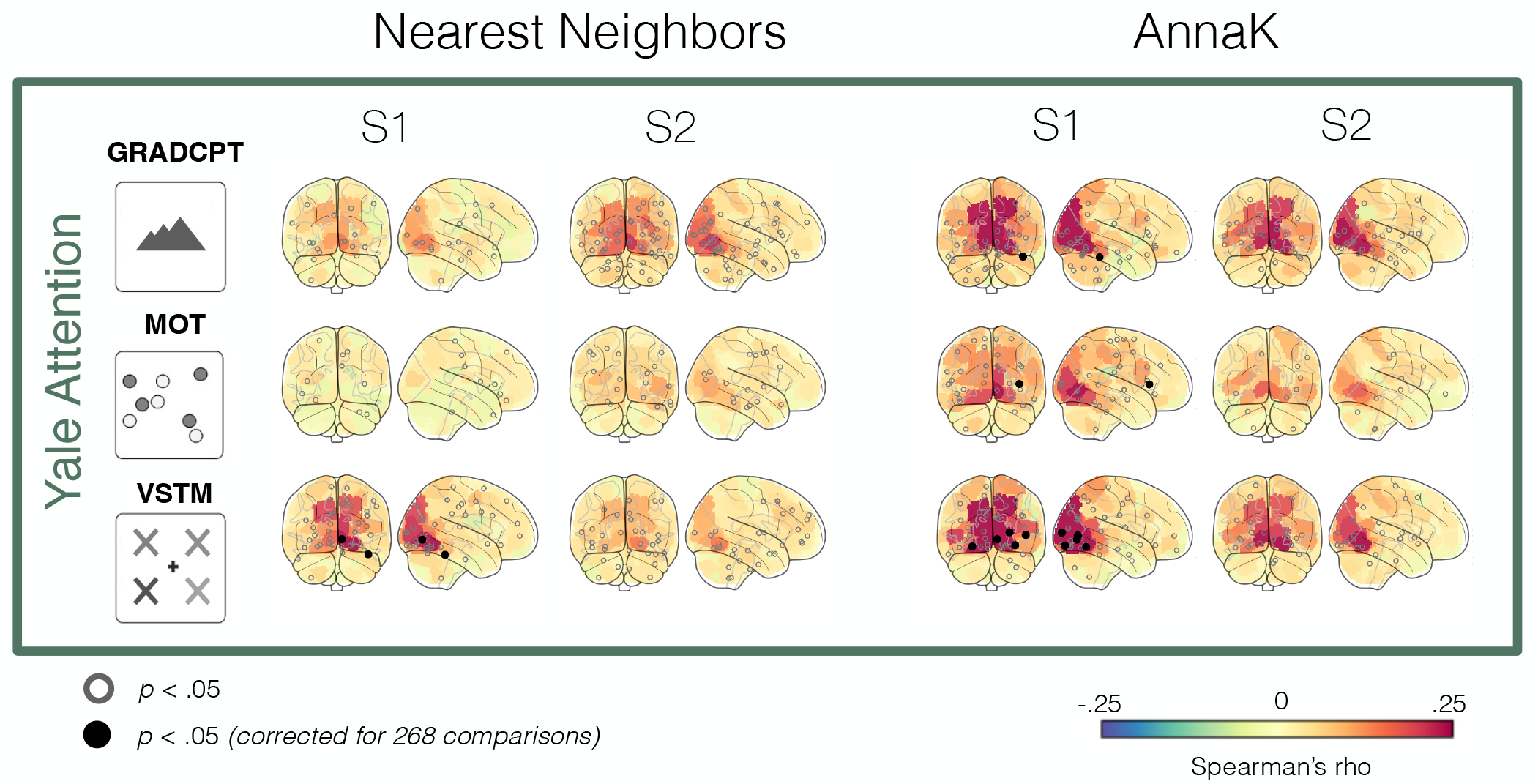
Intersubject RSA (Yale Attention). IS-RSA results for each variable in Yale Attention dataset, separated by movie session (S1 is the first viewing of the film, and S2 is the second). Plotting conventions are the same as figures in the main text (see Figure 3).

**S8:**
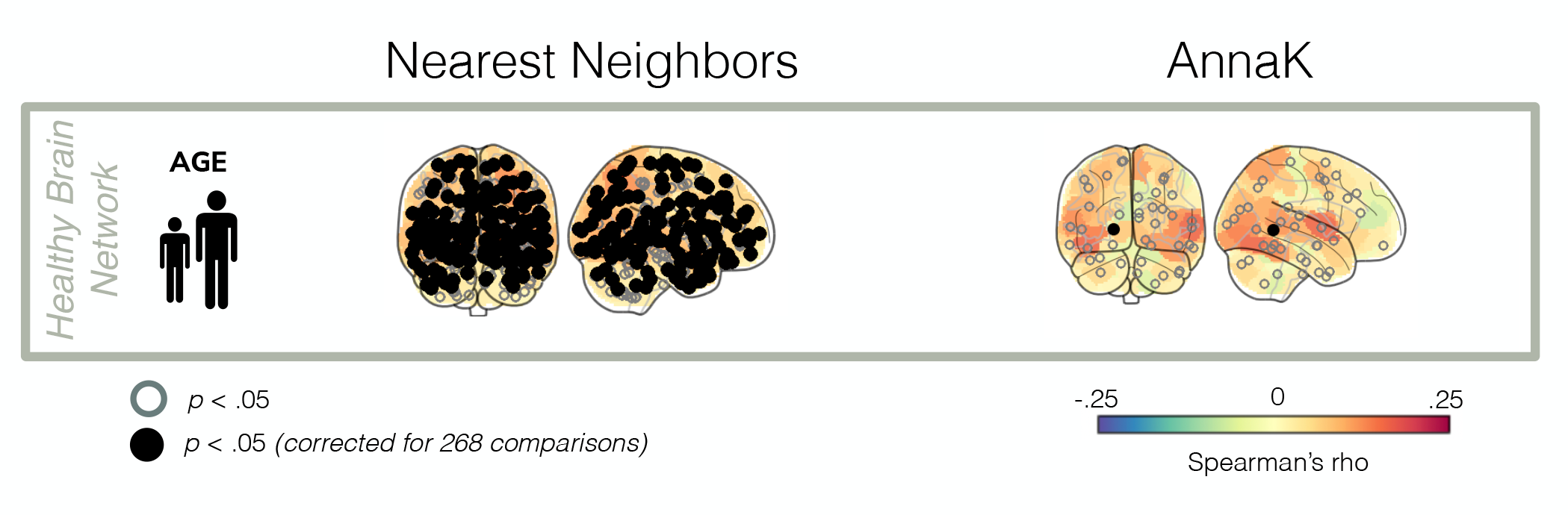
Age Intersubject RSA excluding ages 16 and above. IS-RSA results for age in the Healthy Brain Network sample, but excluding participants aged 16 years and older (the main text includes all available ages, 6-21). Plotting conventions are the same as figures in the main text (see Figure 3).

